# The risk of sexual reproduction promotes the evolution of regulation between host and symbionts

**DOI:** 10.1101/2025.07.30.667627

**Authors:** Alkmini Zania, Paulien Hogeweg, Sam von der Dunk

## Abstract

Sexual reproduction is a widely spread feature of eukaryotes and was already present in the last eukaryotic common ancestor (LECA). Most extant eukaryotes inherit mitochondria from a single parent, but the mechanisms enforcing uniparental inheritance vary widely. Yet, because the first eukaryotes would not have evolved such mechanisms, sexual cell fusion would have inherently led to mitochondrial mixing. Here, we explore the evolutionary consequences of biparental inheritance of endosymbionts during host–symbiont co-evolution using a multilevel, individual-based model of endosymbiosis. Our results show that biparental inheritance introduces evolutionary conflict, as it facilitates the spread of fast-replicating symbionts, which can drive host populations to extinction. However, in a diverse environment, holobionts diversify and adapt to distinct niches, protecting the population from total collapse caused by selfish symbionts. Moreover, this conflict can be resolved through the evolution of signaling mechanisms that allow hosts to regulate symbiont cell cycles. In many cases, sexually reproducing populations not only survive but also outperform their asexual counterparts. We conclude that sexual reproduction could have appeared early during eukaryogenesis.

## Introduction

The emergence of eukaryotes marked a major evolutionary transition, constituting the conversion of a partnership between an archaeal host and an alphaproteobacterial endosymbiont towards a eukaryotic cell [1–3]. The last eukaryotic common ancestor (LECA) possessed hallmark features of eukaryotes such as a nucleus, dynamic cytoskeleton, mitochondria, and meiosis, but the timing and sequence of these innovations remain unresolved [4, 5]. Among them, mitochondrial acquisition was especially transformative, providing bioenergetic advantages [7], enabling genome expansion via endosymbiotic gene transfer [6], and potentially triggering adaptations such as nuclear compartmentalization and sexual reproduction in response to introns and reactive oxygen species [8].

As the endosymbiont transitioned into a mitochondrion, its genome was dramatically reduced — a pattern commonly seen in symbiotic relationships [26]. Many of the proto-mitochondrion’s genes were either transferred to the host nucleus or replaced by host-encoded proteins [27]. Regulatory control by the host evolved through the development of signal peptides and a protein import machinery that was likely co-opted from the symbiont’s protein export systems [28]. These developments were central to resolving conflicts between selection at the host-level and selection at the symbiont-level. [24].

One challenge inherent to mitochondrial endosymbiosis under sexual reproduction is the distribution of mitochondria to daughter cells. In most extant eukaryotes, mitochondria are inherited from a single parent, a pattern known as uniparental inheritance. Uniparental inheritance reduces horizontal mixing among symbionts, possibly suppressing selfish mitochondrial variants [10], maintaining mitonuclear co-adaptation [11], and limiting harmful heteroplasmy [12, 13]. However, uniparental inheritance also increases susceptibility to mutation accumulation via Muller’s ratchet [9]. Despite its wide spread, uniparental inheritance is not universal: instances of biparental inheritance have been observed across fungi, animals, and plants [15, 16], and enforcement mechanisms vary widely among lineages [17, 18]. These patterns suggest frequent evolutionary turnover, shaped by competing selective pressures. For instance, the constant appearance of selfish mitochondrial variants may drive selection for uniparental inheritance, whereas the accumulation of mitochondrial mutations through Muller’s ratchet could provide selection for biparental inheritance [18].

While progress has been made in understanding the evolution of mitochondrial inheritance in extant eukaryotes, its origin during eukaryogenesis remains unknown. Two previous studies proposed that uniparental inheritance had to emerge early during eukaryogenesis, based on its large benefits mentioned before [20, 21]. Yet, proto-eukaryotes may have initially lacked the cellular machinery for selective inheritance, and thus were restricted to biparental inheritance [22].

Previous theoretical models have examined whether the evolution of sexual cell fusion could have been driven by the benefits of mixing symbionts; either as a means of promoting proto-mitochondrial complementation [23] or because of the short-term advantage of homogenizing the cytoplasm when there are negative epistatic interactions among mitochondria and weak mito-nuclear associations [14]. However, another model [24] suggests that in order for cell–cell fusion to evolve, the new holobiont should have already evolved conflict mediation mechanisms, like signaling and transfer of mitochondrial genes to the nucleus, to prevent the spread of selfish symbionts. Therefore, they propose that sex appeared in the later stages of eukaryogenesis. Here, we use a model with more degrees of freedom to explore the dynamics of endosymbiosis under biparental inheritance. Entities are individually simulated, and their growth dynamics arise from their genome. Host and symbiont genomes can evolve complex cell cycles, allowing behaviors such as selfishness, cooperation, and cell-cycle coordination to emerge. We focus on early eukaryotes, assuming no machinery for selective mitochondrial transmission — so fusion leads to biparental inheritance. We then ask: *Under what conditions can holobionts survive despite evolutionary conflict? And what mechanisms might evolve to mitigate this conflict?*

## Methods

### Model of cell cycle regulation

Our modeling framework is based on the multilevel computational model introduced by von der Dunk et al. [29] to simulate the evolution of cell cycle regulation. Each organism encodes a Boolean gene regulatory network on a linear, beads-on-a-string genome. The genome contains regulatory genes and binding sites, creating expression dynamics that result in a cell cycle. Progression through the four main stages (G1, S, G2, M) depends on the expression of five core regulatory genes (g1–g5). Regulation is stochastic: gene products bind upstream binding sites probabilistically based on bitstring similarity and contribute to activation depending on site- and gene-specific weights and thresholds. The genome also contains household genes that are not active, but have to be carried by the organism in order for it to be viable.

Genome replication depends on nutrient availability: during each timestep in S-phase, a cell uses nutrients in its local environment to replicate genome beads (binding sites, regulatory genes, or household genes). If a cell attempts to divide before completing genome replication, it dies. When division is successful, a daughter cell is created in a random adjacent grid site, replacing any previous occupant of that site. The model is set on a grid with uniform nutrient influx (50×50, *n*_*influx*_ =30) or on a nutrient gradient consisting of 11 sectors from nutrient-rich to nutrient-poor (25×275, 80 > *n*_*influx*_ > 2).

Two distinct adaptive strategies can be distinguished among populations of cells evolved on the nutrient gradient: specialists and generalists [29]. Specialists evolved to regulate an S-phase of fixed duration, locally adapted to the nutrient influx of the sector they inhabit. In contrast, generalists tune their cell cycle duration according to nutrient presence.

### Endosymbiosis

The model of Von der Dunk et al. (2023) was extended [19, 25] to study obligate endosymbiosis. Now, each cell, which we call ‘holobiont’, consists of one host and one or more symbionts. Host and symbiont genomes evolve independently but interact through the shared nutrient environment and, when enabled, molecular communication. Passive communication proceeds through leakage of expressed regulatory product from host to symbiont or from symbiont to host with a probability of 1% per product. Active communication proceeds through the relocalization between compartments of gene products as defined by their signal peptides. Products that are leaked or transported from host to symbiont and vice versa function as native products. Holobionts reproduce when the host divides; symbionts are distributed between parent and offspring stochastically. Holobionts without symbionts die.

### Biparental inheritance of symbionts

The model was extended with sexual reproduction of holobionts, implemented as biparental inheritance of symbionts. Hosts ready to divide seek a mating partner within the same nutrient sector - another cell ready to divide. If no partner is found in a timestep, the holobiont pauses its cell cycle until a match appears. Mating results in two offspring holobionts; each one receives the full host genome of one parent and inhabits a cell adjacent to that parent. Symbionts from both parents are randomly distributed between the two daughter cells.

### Experimental setup

Initial holobiont species were formed by pairing pre-evolved hosts and symbionts (species 5-11, Table 1, Table S1), each adapted as free-living individuals on the nutrient gradient. These evolved species are described in [29]. We chose strains R2 (a specialist optimized for rich-nutrient conditions) and R3, R8, and R9 (generalists that can survive in a broader range of nutrient conditions). Each species then evolved under homogeneous intermediate conditions (*n*_*influx*_ = 30) or on a nutrient gradient independently (Table 1). Experiments on the homogeneous grid took place under different host–symbiont communication modes enabled; no leakage nor signaling, only signaling, and signaling with leakage. Signaling is not present in initial holobionts but is allowed to evolve through mutations of the signal peptide, which is present in all regulatory genes.

**Table 1.**
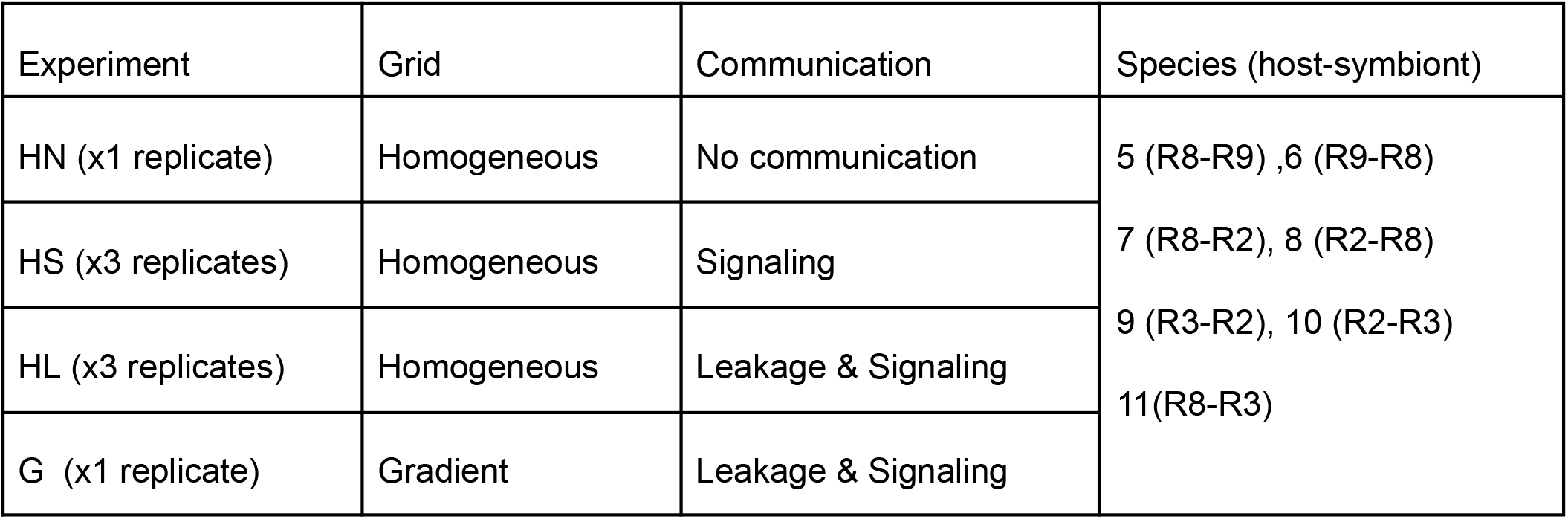
Overview of experiments.

#### Box 1. Summary of previous results

**Primitive strains**: Identical host and symbiont that can perform a rudimentary cell cycle.

**Pre-evolved strains**: Host and symbiont initially evolved independently as free-living cells on the nutrient gradient and can perform complex cell cycles. Holobionts were made by pairing different strains in host - symbiont relationships.

Pre-evolved strains and primitive strains without product leakage evolved implicit cell cycle co-ordination (fig B1). Host and symbiont do not evolve signaling, but specialize on different nutrient concentrations, forming a stable equilibrium in their growth rates.

**Figure B1.**
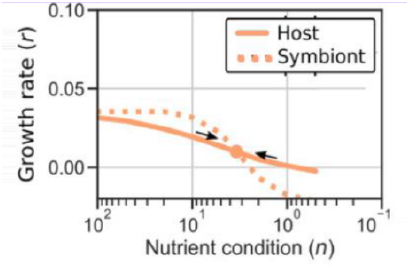
Implicit control. The growth curves of host and symbiont are constructed by assesing their growth rates independently in different nutrient conditions. In implicit co-ordination, they form a stable equilibrium at the nutrient concentration for which their growth rates are equal. Figure from von der Dunk et al., 2023.

In contrast, primitive holobionts that evolved with passive product leakage and the ability to signal between hosts and symbionts evolved three distinct mechanisms of explicit cell cycle coordination (fig. B2):

a) **Host control**. The host controls its own cell cycle as well as the symbiont’s. This is achieved through dual localization of host genes, sometimes leading to synchronization of their expression dynamics.
b) **Bi-directional control**. Both symbiont and host influence each other’s cell cycle through signaling, establishing check points for symbiont number and host genome replication to ensure safe division.
c) **Symbiont control**. A rare case of signaling where only the symbiont signals to the host, informing it that there are enough symbionts to safely divide.

**Figure B2.**
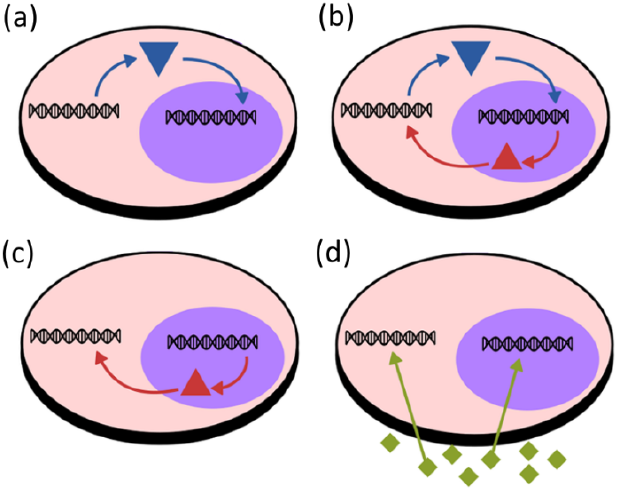
Four unique evolved mechanisms of cell-cycle coordination: (a) host control, (b) bi-directional control, (c) symbiont control, and (d) implicit control. Gene products can be transported from one compartment to another and bind to promoter sites there. Figure adapted from von der Dunk et al., 2024.

## Results

### Selfish symbionts emerge, but can be controlled through signaling

We investigate the effects of sexual reproduction with biparental inheritance on the evolution of endosymbiosis between complex, distantly related organisms. In earlier work, pre-evolved ‘prokaryotes’ were paired as hosts and symbionts (table 1) and evolved reproducing clonally [19]. These experiments showed that clonal holobionts did not evolve signaling, though mutations that would allow it were included (see Appendix for one exception). Instead, they evolved implicit cell-cycle coordination by forming growth curves with a stable equilibrium [19] (box 1). In contrast, when the same experiments are repeated under biparental inheritance, selfish symbionts that outgrow their host emerge, destabilising the holobiont.

Populations that evolved on the homogeneous grid often collapsed due to the uncontrolled replication of selfish symbionts, especially without communication (fig. 1a). In rare instances, hosts survived despite extreme symbiont loads depleting nutrients to a bare minimum level resulting in very slow growth. Only one host – symbiont pair (species 9) invariably evolved a stable population in all conditions (fig. 1a-c), with a large symbiont genome and a small host genome.

**Figure 1.**
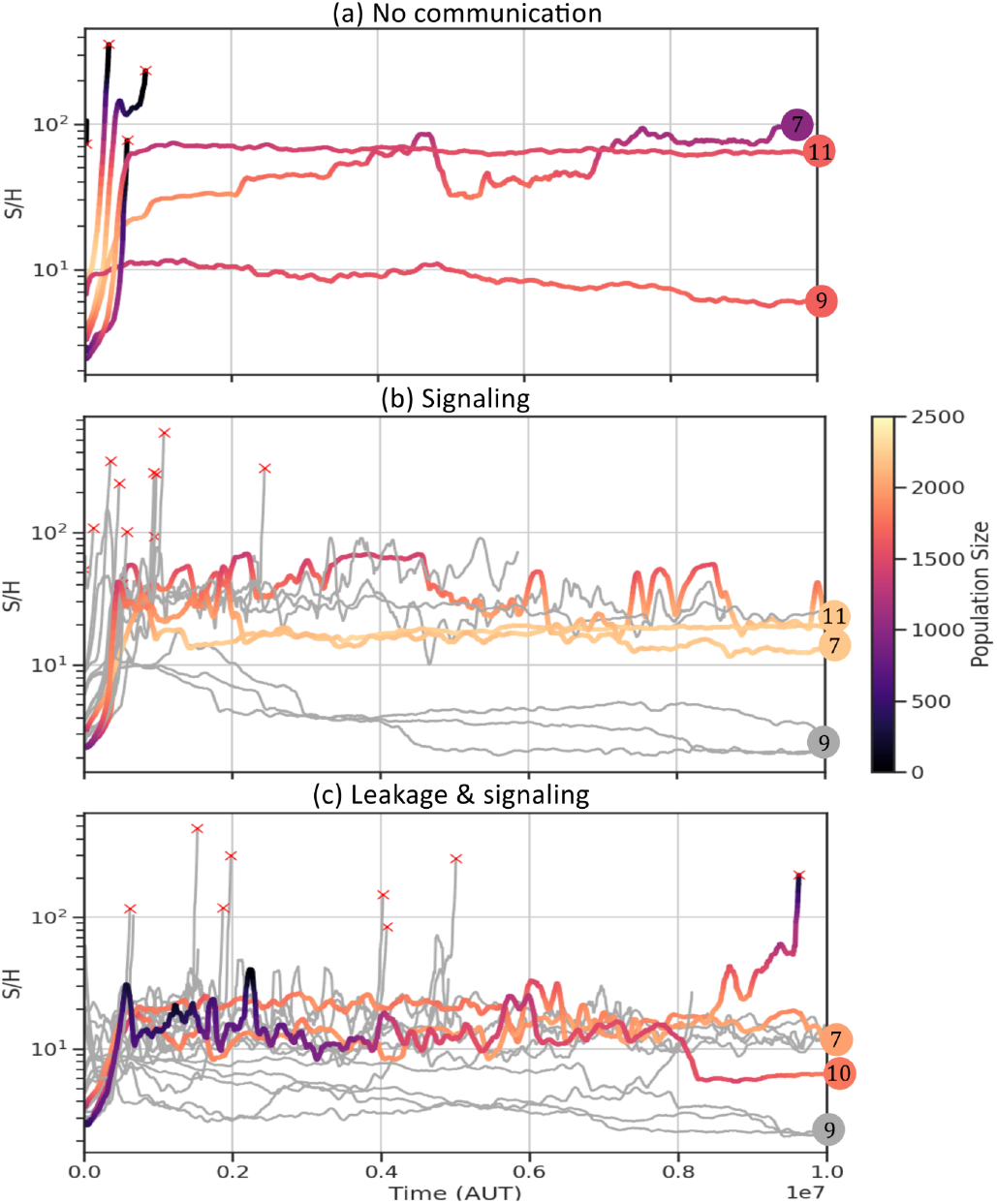
Symbiont numbers fluctuate in populations evolving on homogeneous nutrient influx (H). (a) Populations with no host – symbiont communication (b) with signaling (c) with leakage and signaling. Each line represents a distinct population. Symbiont per host scale is logarithmic. Numbers in circles indicate the starting species of selected populations. Populations with signaling (HS) and leakage & signaling (HL) were initiated 3 times for each species. Symbiont numbers and host population size fluctuate highlighting the intragenomic conflict. Many populations go extinct (indicated by a red ‘x’). Without communication (HN) only 3 populations survive; HN9, that never exhibited strong host symbiont conflict, and HN7 and HN11, with extreme symbiont numbers. Species 7 and 11 were also more likely to survive when communication between compartments is possible, but with better controlled symbionts. With more (possible) routes of communication, more populations survive and tighter control of symbionts is achieved.

Horizontal mixing provides an advantage to fast-replicating symbionts, who will spread over the population, instead of being confined to one lineage. Selection for fast replication favours symbionts with small genomes and few household genes, enabling rapid replication even under limited nutrient conditions. Over time, these selfish symbionts proliferate and deplete resources for the host to the point of starvation.

While holobionts evolving on the homogeneous grid without host-symbiont communication (i.e. leakage and signaling) were highly susceptible to extinction, populations with leakage or the potential for signaling demonstrated longer survival times and lower symbiont numbers (fig. 1). Some populations not only avoided extinction but also stabilized symbiont numbers through the evolution of signaling. In previous clonal experiments [25] leakage was deleterious for holobionts, who suffered a drop in population size and symbiont numbers, even after long periods of evolution. However, under biparental inheritance, the reduction of symbiont numbers due to leakage increases the survival of holobionts.

Despite the existence of conflict resolution strategies, stabilization was not universal. In many surviving populations, host-symbiont conflict persisted, manifesting as fluctuations in host and symbiont numbers over time. These instabilities reflect the ongoing evolutionary battle between host control and symbiont selfishness.

Evolution of signaling in pre-evolved holobionts was a surprising outcome, because under clonal reproduction pre-evolved holobionts never evolved any form of explicit communication (even when host and symbionts were identical) [25]. The evolution of signaling may be particularly challenging between unrelated pre-evolved organisms, where regulatory networks and binding site sequences have already diverged significantly. In such cases, achieving functional host-symbiont communication requires overcoming genetic and regulatory incompatibilities, making signaling a less accessible solution compared to implicit coordination. However, we think the host—symbiont conflict introduced by biparental inheritance increases selection for conflict resolution and overcomes these regulatory challenges. Evolution of signaling allowed hosts to interfere with, regulate, or take control of symbiont cell cycles.

### Diversification on the nutrient gradient protects from extinction

In contrast to the homogeneous grid, populations that evolved on the nutrient gradient did not go extinct within the timeframe of the experiments. The gradient environment induced diversification of holobionts (fig. 2d). While selfish symbionts frequently emerged, their effects were localized, devastating strains of a few sectors rather than the whole population (fig. 2a, b). Host signaling also evolved in populations on the gradient, but not as frequently as it did on the homogeneous grid. In population G8 (fig. 2), signaling evolved, but host and symbiont retained regulatory autonomy and exhibited implicit control (fig. 2c).

**Figure 2.**
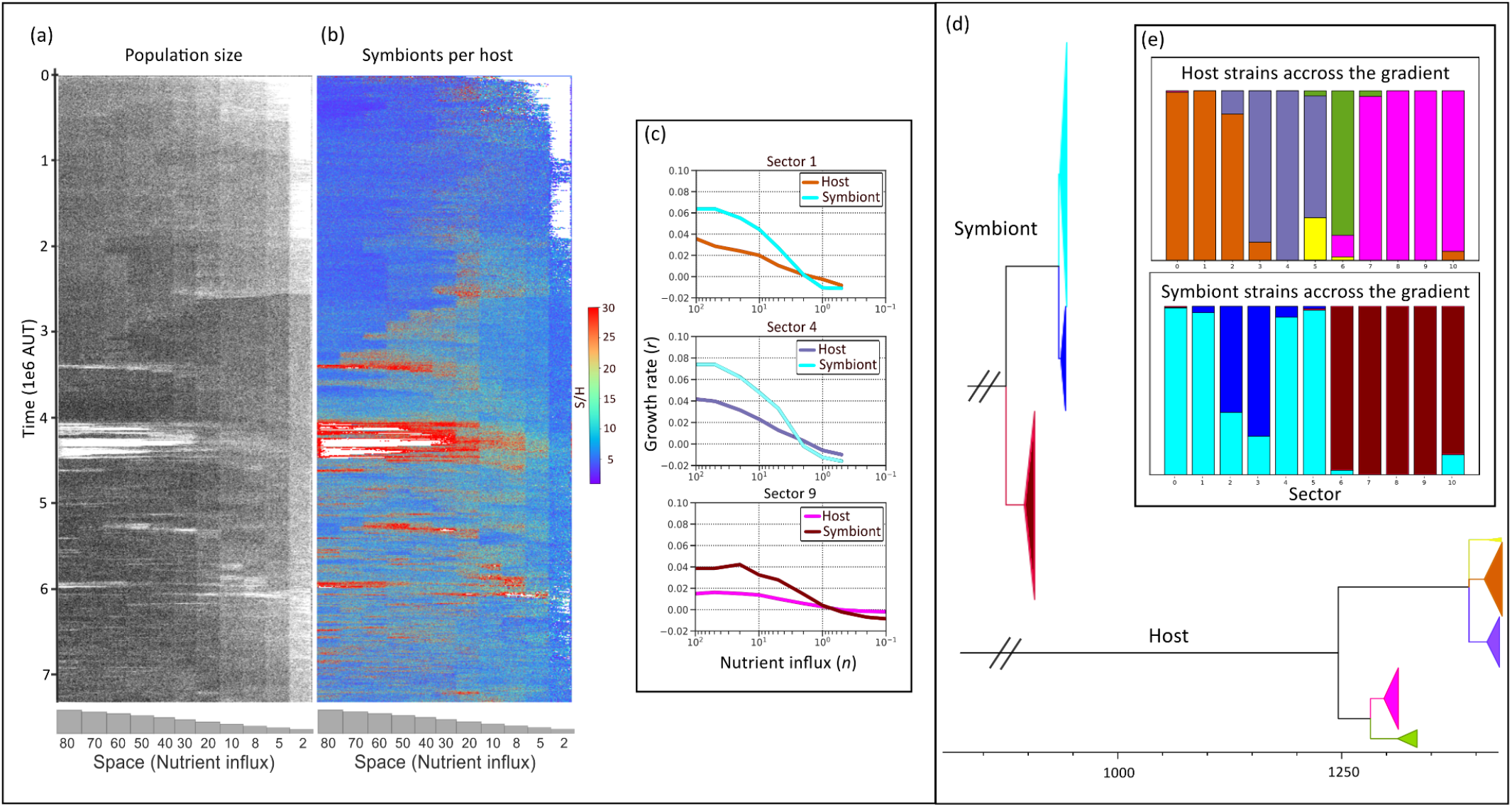
Species 8 evolving on the nutrient gradient under biparental inheritance. (a) Spacetime plot showing the size of population G8 evolving on the nutrient gradient. Darker shade indicates a larger host population. (b) Spacetime plot showing symbionts per host numbers in population G8. Red color indicates high symbiont numbers. White indicates empty space. Selfish symbionts invade and often kill hosts in different sectors of the gradient, most noticeably around 4.5M AUT. Annihilated sectors are then reinvaded by neighboring hosts. (b) Growth curves (see fig. B1) of most abundant hosts and symbionts in different sectors of the gradient. Strains in nutrient poor sectors grow slower but survive in harsher conditions compared to strains in richer sectors. Growth curves are colored according to the clade of (c) to which they belong (c) Phylogenetic tree of hosts (bottom) and symbionts (top), colored by strains that inhabit different sectors of the gradient. (d) Presence of host and symbiont strains on the nutrient gradient at 7M AUT.

One potential reason for the increased population stability on the gradient is that the diversification across the grid creates host – symbiont incompatibilities. Thus an emerging selfish symbiont strain is not easily able to invade all sectors (fig. 2b). Another factor that could add to the evolutionary resilience of populations on the gradient is the fact that mating only takes place locally, thereby imposing a delay on the invasion of symbionts into adjacent sectors.

Another interesting observation is that on the gradient, holobionts with leakage and signaling achieved larger population sizes under biparental inheritance, compared to their clonal counterparts, even under stress from selfish symbionts (fig. 3). In contrast to clonal populations, where leakage was harmful, under biparental inheritance the extra stress of leakage can improve survival and increase population size, in line with observations on the homogeneous grid.

**Figure 3.**
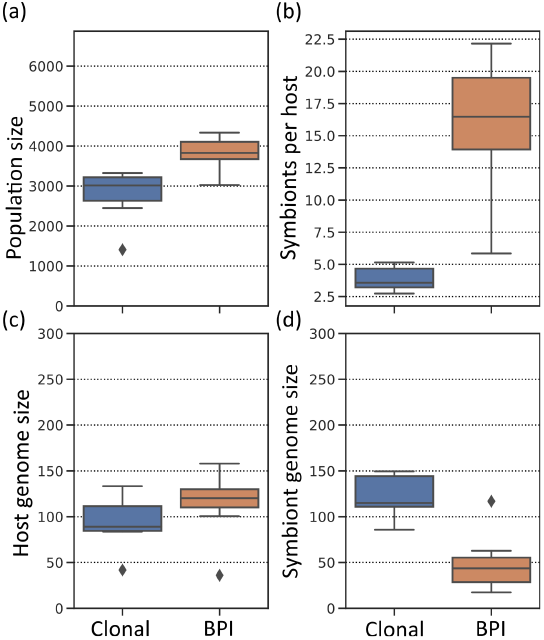
Overview of holobiont populations evolved with leakage and signaling on the nutrient gradient at 4 × 10^6^ AUT. In populations with biparental inheritance (BPI) (a) population sizes (b) symbiont numbers (c) host genome size are increased, and (d) symbiont genome size is decreased compared to clonal populations. Increased symbiont numbers reflect ongoing conflict and reduced integration.

### Evolution of genome sizes

Competition among symbionts for rapid replication drove a consistent reduction in symbiont genome size (fig. 4). This reduction was predominantly achieved through the loss of household genes, concurrent with gain of household genes in the host. As these genes are essential for the holobiont’s survival, an asymmetry evolved between the two compartments: hosts bear the cost of household genes while symbionts streamlined their genomes to optimize replication speed. In some populations not all symbiont household genes were lost in the timeframe of the experiments (4M AUT), but the symbiont genome size displayed a continuous decline. Species 9, which is the only species that survived biparental inheritance in all experimental settings and replicates, stands out of this pattern by transferring most household genes to its symbiont. This reverse genome size asymmetry highlights that the symbionts of species 9 were not prone to the evolution of selfishness.

**Figure 4.**
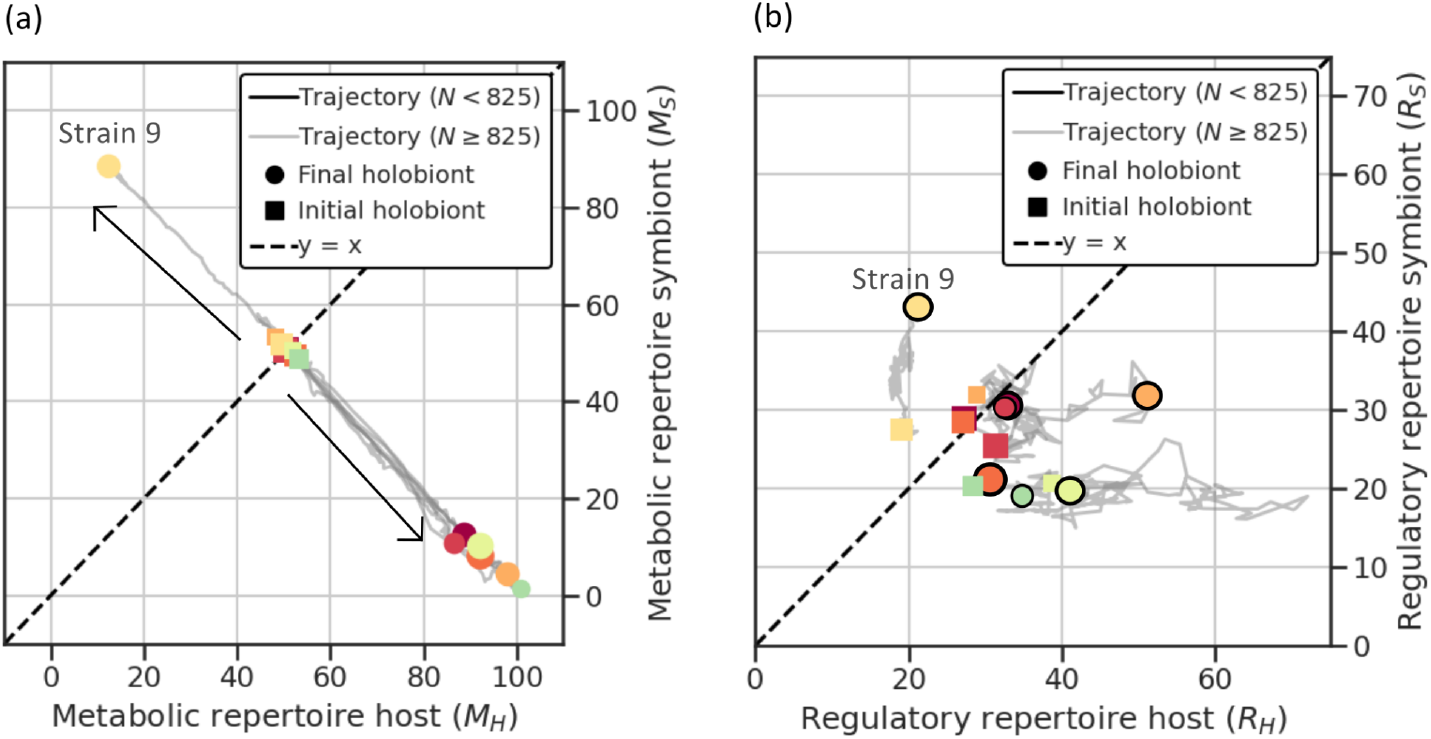
Host and symbiont genome composition after evolution on the gradient. (a) Household gene distribution between hosts and symbionts across populations evolved on the nutrient gradient. Most populations show a near-complete transfer of household genes to the host, with symbionts retaining minimal household gene content. (b) Regulatory gene distribution between hosts and symbionts. Symbionts generally retain fewer regulatory genes than their hosts, reflecting their reduced genome size. The exception is Population G9, where the symbiont genome retains a significant number of regulatory and household genes, diverging from the general pattern. Each data point represents a distinct holobiont population at 4M AUT.

Even regulatory genes are impacted by the strong selection pressure on genome size in hosts and symbionts. In most species, the regulatory repertoire of the host is larger than that of the symbiont, despite fluctuations along evolutionary trajectories and the absence of a clear tradeoff between host and symbiont repertoires.

Since replication speed depends strongly on genome size, smaller genomes confer a selective advantage by allowing symbionts to complete replication and division more quickly. This consistent symbiont genome streamlining is in contrast to clonal experiments, where there was no strong selection for small symbionts [19]. However, clonal strains that did evolve smaller symbionts, were more successful in terms of population size.

### Types and function of signaling

Biparental inheritance introduced strong intragenomic conflict between symbiont and host, resulting in extreme fluctuations in evolutionary time, and often extinction. Under biparental inheritance, signaling in holobionts is used as a means to control symbionts or even resolve the conflict and stabilize populations. Across experiments, different strategies evolved, with varying degrees of regulatory integration and effectiveness in stabilizing symbiont numbers. below some of these evolved regulatory mechanisms are examined in detail.

Cell-cycle synchronization was the most effective strategy for stabilizing host—symbiont interactions, as seen by the tight control of symbiont numbers over evolutionary timespans (fig. 5a, observed at ∼8M AUT). Cell-cycle synchronization is achieved through the complete takeover of the symbiont cell cycle by the host. In HL10, only two strong regulatory interactions between genes are still encoded by the symbiont (fig. 5c). Thus, symbionts have lost their autonomy and cannot become selfish by outgrowing their host. The extensive regulatory integration of host and symbiont resolves the intragenomic conflict, yielding large population size and stable symbiont numbers (fig. 5a). Interestingly, synchronization drives the evolution towards equal genome sizes of host and symbiont, contrasting with the general trend of genome size asymmetry (previous section). In fact, HL10 exhibited asymmetry in genome size for a long time, but once cell-cycle synchronization was achieved, this asymmetry gradually disappeared (fig. 5b); as division happens simultaneously, it is favorable for the holobiont to finish genome replication in all compartments at the same time. The long term stability of populations with a synchronized cell cycle is also observed in populations that started with identical host and symbiont, where signaling is easier to evolve due to smaller genetic differences between the two compartments (Appendix).

**Figure 5.**
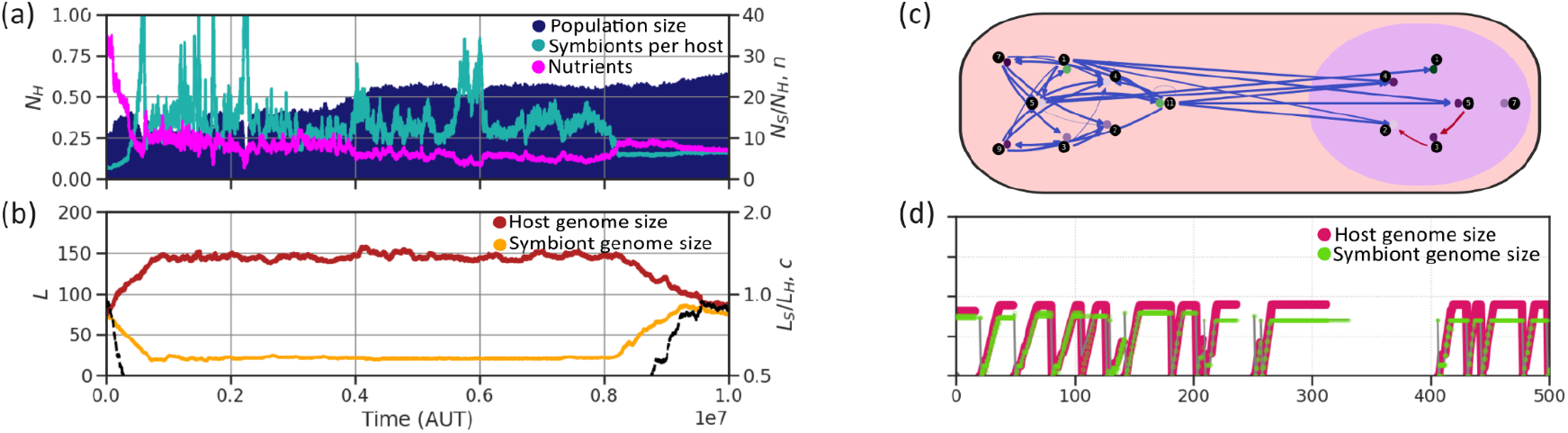
Evolution of population HL10c to a synchronized cell cycle. (a) Timeline of the population evolving on constant nutrient influx n_influx_ = 30, with leakage & signaling. After 8M AUT of host – symbiont conflict, the endosymbiotic relationship is stabilized. (b) Timeline of the evolution of host and symbiont genome sizes. Before synchronization at 8M AUT, the symbiont’s genome is streamlined and the host’s genome increases by carrying household genes. After synchronizations the two compartments evolve equal genome size. (c) Regulatory networks that give rise to a host controlled synchronized cell cycle. Interactions show the aggregate regulatory effect (binding probability times regulatory weight) of gene types (black disks) on loci (colored disks adjacent to the gene types). Arrow thickness shows probability of interaction and color compartment of origin (red: symbiont, blue: host). Genes are colored based on their activation threshold (green < 0, purple > 0). (d) Cell cycle progression of HL10c in host and symbiont: measured by how much of the genome has been replicated (genome size can be read off from the maxima). Genome replication and division happen simultaneously for host and symbionts at 10M AUT.

*Host control of symbiont cell cycle progression* was another strategy that effectively stabilized intragenomic conflict. In HS7, holobionts achieved partial cell-cycle synchronization while maintaining relatively high symbiont numbers (fig. 6a). Unlike for full cell-cycle synchronization, host and symbiont retained regulatory autonomy (fig. 6g), allowing independent control of their respective cell cycles. Yet, the emergence of fast-growing (selfish) symbionts is prevented by the combination of growth and regulatory dynamics in this population. The host functions as a slow-growing generalist, while the symbiont is a fast-growing specialist that thrives only in nutrient-rich environments, as witnessed by their growth curves (fig. 6g). However, the host delays symbiont division while remaining in the S phase, enabling symbiont survival under nutrient-poor conditions. As a result, signaling increases the replication of symbionts, who survive in harsher conditions and achieve larger numbers (fig. 6f). At the same time, it stabilizes their relationship with the host by preventing unchecked proliferation, resulting in a partially synchronized cell cycle (fig. 6d).

**Figure 6.**
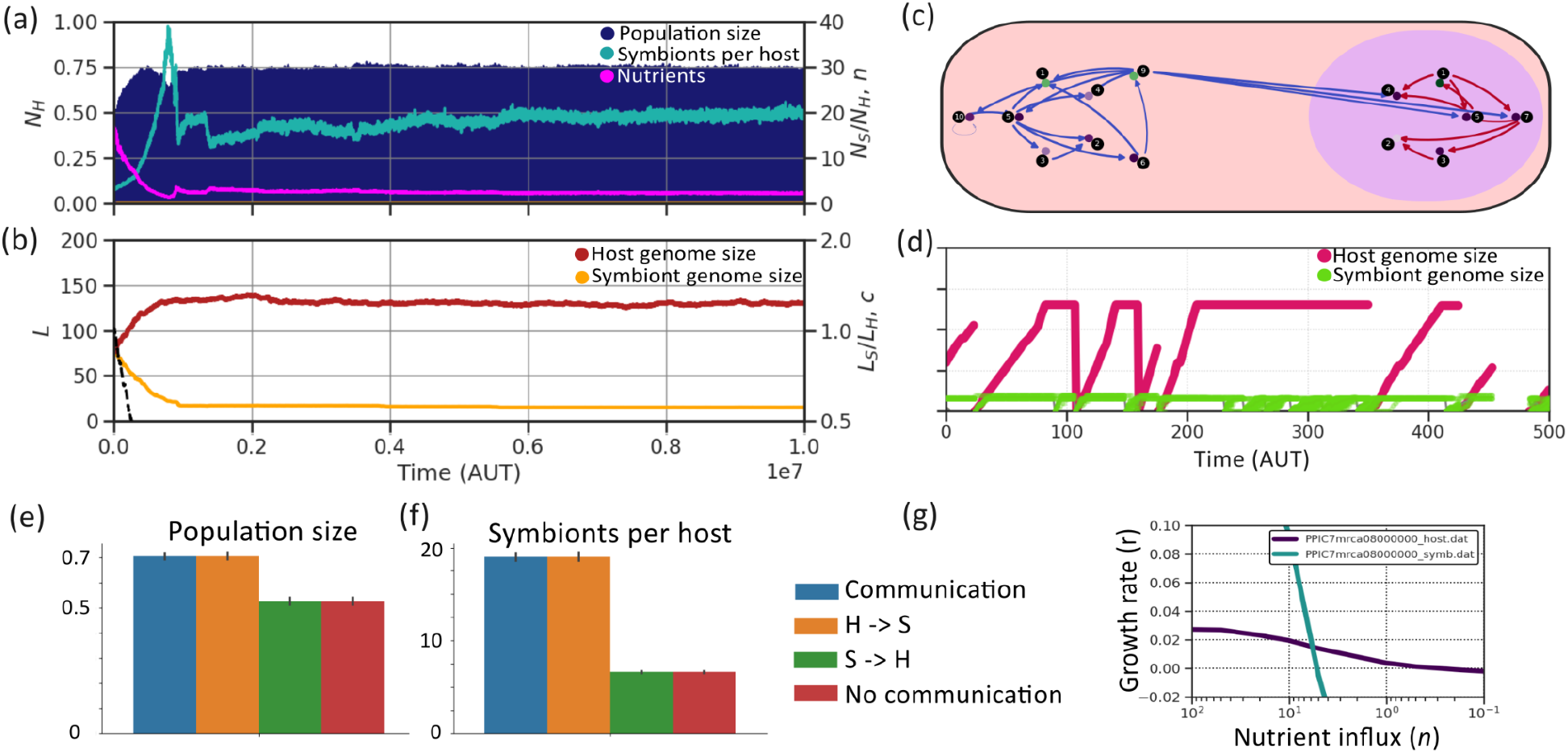
Evolution of population HS7c to a partially synchronized cell cycle. (a) Timeline of population HS7c evolving on constant nutrient influx n_influx_ = 30, with signaling but no regulatory product leakage. After a short explosion in symbiont numbers around 1M AUT, the host population stabilizes and maintains stable, though high, numbers of symbionts. (b) Timeline of the evolution of host and symbiont genome sizes. The symbiont is streamlined and the host carries most household genes. (c) Gene regulatory network of host and symbiont MRCA at 10M AUT. While the host remains in the S-phase, it activates the symbiont’s S-phase genes too. (d) Cell cycle progression of host and symbiont: measured by how much of the genome has been replicated. Both compartments replicate their genome at the same time and the symbiont stalls its division, but often divides before the host. (e) Population size and (f) symbionts per host under different communication conditions. Host signaling allows symbionts to survive in low nutrients, by keeping them in the S-phase long enough to replicate their genome. Without host signaling, the population survives with implicit coordination, but with smaller population size. (g) Growth curves of host and symbiont MRCA (t=8M AUT). Both host and symbiont remain autonomous. The host is a slow generalist and the symbiont a fast specialist that only survives in rich environments. The assessment of symbionts per host and population size is performed on the most recent common ancestor (MRCA) of each population, without mutations and under homogeneous nutrient influx.

*Host interference in symbiont’s cell cycle* was a less effective but more easy to evolve strategy, whereby hosts delay the symbionts’ cell cycle. Signaling is often weak and unstable over time, as symbionts that escape host control can proliferate more rapidly. Despite regulatory interactions between host and symbiont, both compartments remain fully autonomous and can survive independently of signaling (fig. 6c, g). In both populations, the symbiont’s genome remains compact, with few or no household genes, favoring rapid growth.

In one replicate of population HS11 (fig. 7a-d), weak host-derived signals (fig. 7b) prevent the symbiont from completing its cell cycle while the host remains in the S-phase. Host signaling appeared at 2.2M AUT and progressively strengthened over time, leading to a gradual reduction in host-symbiont conflict. However, fluctuations in symbiont numbers indicate that the conflict remains unresolved (fig. 7a).

**Figure 7.**
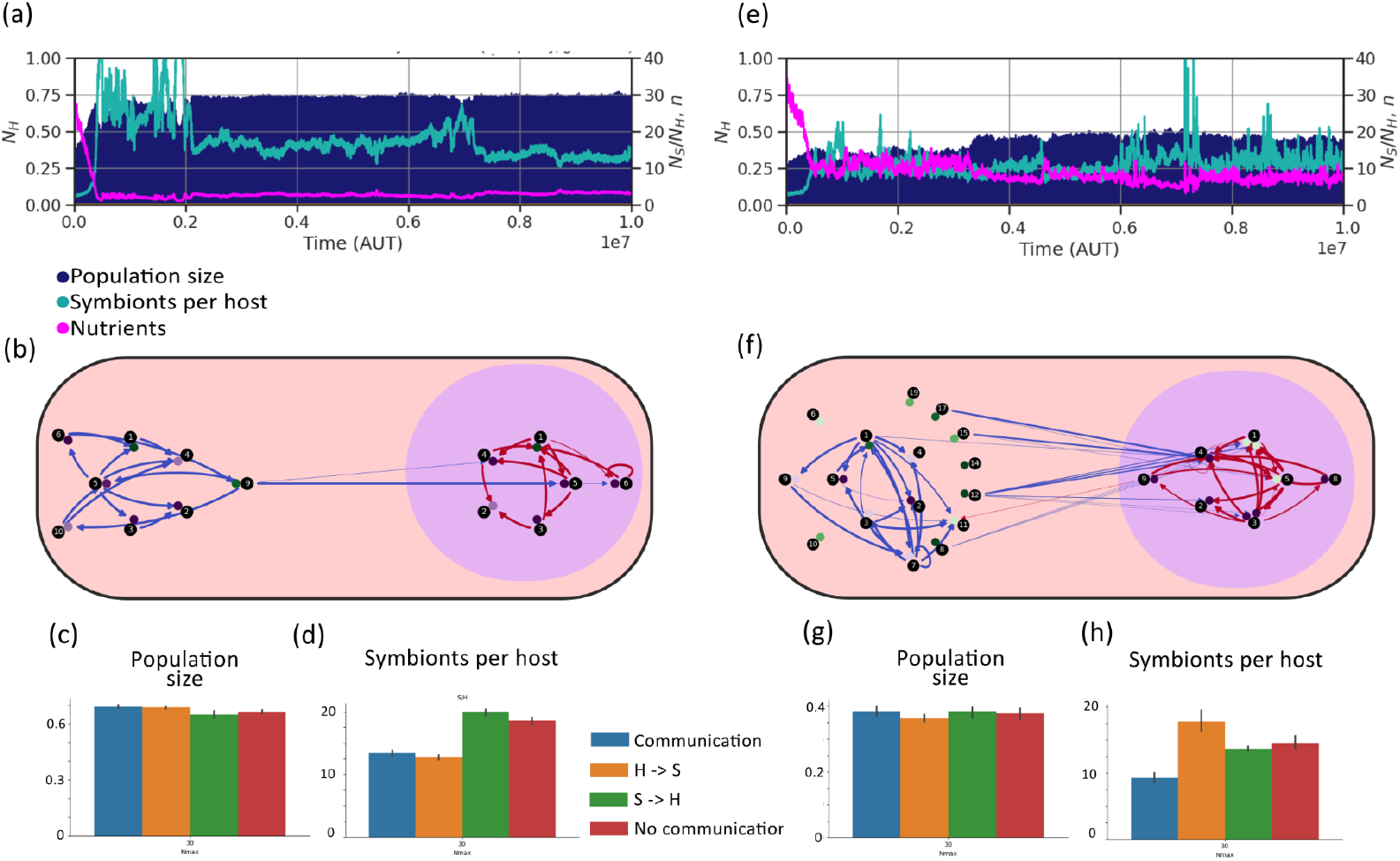
Evolution of populations HS11c and HL10b, under host – symbiont conflict. (a, e) Timeline of (a) HS11c (no leakage, transfer) and (d) HL10b (leakage & transfer) on constant nutrient influx n_influx_ = 30. Host – symbiont conflict is present, as symbiont numbers fluctuate sometimes decreasing host population size. (b, f) Gene regulatory network of HS11c (b) and HL10b (f), which have evolved host controlled and bidirectionally controlled signaling accordingly. (c, g) Population size of HS11c (c) and HL10b (g) under different communication conditions (see fig. 6). (d, h) Symbionts per host in HS11c (d) and HL10b (h) under different communication conditions. Leakage background is the same as in original populations: off for HS11c on for HL10b. Full signaling decreases symbiont numbers in these populations.

In one replicate of population HL10 (fig. 7e-h), the host constitutively expressed genes that target symbiont regulatory pathways (fig. 7f), potentially interfering with the symbiont’s cell cycle. When signaling is blocked, the holobiont maintains a larger symbiont population. However, host-derived signaling, along with symbiont feedback, is required to regulate symbiont numbers effectively (fig 7h).

## Discussion

In this study we have explored the implications of biparental inheritance of symbionts, using an evolvable system with a complex genotype-phenotype map. We modeled pre-evolved hosts and symbionts individually, allowing selfishness, co-operation and regulatory integration of initially autonomous replicating entities to emerge.

Consistent with early hypothesis, biparental inheritance of endosymbionts introduces an evolutionary conflict between host and symbiont genomes. In contrast to clonal reproduction, where symbionts are vertically transmitted within a lineage, horizontal mixing in sexual reproduction allows fast-replicating symbionts to spread in the host population. Thus the equilibrium observed in clonal populations, where host and symbiont growth rates co-adapt through nutrient-mediated feedback, becomes evolutionary unstable and selfish symbionts often drive populations to extinction.

Despite the dangers of symbiont mixing, conflict mediation is possible; populations can survive and even thrive under biparental inheritance. The evolution of explicit signaling mechanisms allows hosts to regulate symbiont replication and restore holobiont stability with varying success across experiments. Cell cycle synchronization seemed to completely resolve the conflict as indicated by stable symbiont numbers. Notably, such signaling did not evolve in pre-evolved holobionts under clonal reproduction [25], showing that intragenomic conflict can be a driving force for regulatory integration of distantly related genomes.

Other factors also contributed to population survival apart from signaling. Spatial diversification on the nutrient gradient helped prevent extinction: selfish symbionts were often confined to specific sectors, while neighboring holobionts could recolonize affected areas. The non-homogeneous environment of the gradient can result in the evolution host – symbiont incompatibilities among sectors. In additional experiments where we allow for reversal to asexual reproduction, holobionts mutated back to asexuality, thus confining selfish symbionts (Appendix). As LECA likely reproduced both sexually and asexually [4], occasional cell fusion with biparental inheritance may have occurred without necessarily dooming the population.

A major axis of host–symbiont conflict in our model is selection on genome size, because replication speed directly depends on genome length. Symbionts evolve smaller genomes by shedding household genes, in this way maximizing their division rate. Hosts compensate by retaining household genes, leading to an asymmetry between the two partners. In clonal populations, genome reduction among symbionts was less consistent, but lineages with smaller symbionts achieved the highest population sizes [25], suggesting that symbiont genome streamlining can improve holobiont fitness. While present-day mitochondria do not experience selection for genome streamlining due to cell-cycle constraints, factors such as more efficient resource allocation may have also played a role in the reduction of symbiotic DNA.

The extent to which host–symbiont conflict poses a threat to a population appears to depend strongly on the properties of the initial host–symbiont pair. Species 9 consistently achieved stability without evolving communication, instead maintaining a large, non-selfish symbiont. Species 7 and 11, although initially unstable, survived without signaling and had a higher chance to control their symbionts, as they lived long enough to be able to evolve signaling. On the contrary, species 5, 6 and 8, whose symbionts are high-quality generalists able to outgrow their host, never survived on the homogeneous grid. These examples suggest that apart from metabolic capacities [30], other characteristics of the founding strains (e.g. being a generalist or specialist) shape the process of eukaryogenesis.

In general, biparental inheritance drives selection for genetic integration by strengthening intragenomic conflict, and thereby increasing the evolutionary potential for novel regulatory strategies. Asexual pre-evolved strains almost always maintain static, signaling-free populations, while sexual populations — even those burdened by selfish symbionts — often evolve larger population sizes and more elaborate regulatory coordination. Our findings suggest that cell fusion with biparental inheritance could have emerged before the full domestication of mitochondria, contrary to earlier suggestions [24] as biparental inheritance fosters regulatory integration between genetically distinct partners providing that the foundations for the evolution of signaling are present. With recent discoveries of Asgardarchaea shedding light on early host lineages, our model offers a valuable theoretical context for future empirical investigations into host–symbiont coevolution.

## Acknowledgements

The authors wish to thank Jan Kees van Amerongen for running the local computer cluster and assisting in technical matters.

